# Evidence for an Integrated Gene Repression Mechanism Based on mRNA Isoform Toggling in Human Cells

**DOI:** 10.1101/264721

**Authors:** Ina Hollerer, Juliet C. Barker, Victoria Jorgensen, Amy Tresenrider, Claire Dugast-Darzacq, Leon Y. Chan, Xavier Darzacq, Robert Tjian, Elçin Ünal, Gloria A. Brar

**Author notes:** Present address: Department of Biology, Massachusetts Institute of Technology, Cambridge, MA, USA, 02139. These authors contributed equally to this work (Juliet C. Barker, Victoria Jorgensen). These authors contributed equally to this work (Elçin Ünal, Gloria A. Brar).

## Abstract

We recently described an unconventional mode of gene regulation in budding yeast by which transcriptional and translational interference were used in an integrated manner to down-regulate protein expression. Developmentally timed transcriptional interference inhibited production of a well translated mRNA isoform and resulted in the production of an mRNA isoform containing inhibitory upstream open reading frames (uORFs) that blocked translation of the ORF. Transcriptional interference and uORF-based translational repression are established mechanisms outside of yeast, but whether this type of *integrated* regulation was conserved was unknown. Here we find that, indeed, a similar type of regulation occurs at the locus for the human oncogene *MDM2*. We observe evidence of transcriptional interference between the two *MDM2* promoters, which produce a poorly translated distal promoter-derived uORF-containing mRNA isoform and a well-translated proximal promoter-derived transcript. Down-regulation of distal promoter activity markedly *up-regulates* proximal promoter-driven expression and results in local reduction of histone H3K36 trimethylation. Moreover, we observe that this transcript toggling between the two *MDM2* isoforms naturally occurs during human embryonic stem cell differentiation programs.

## INTRODUCTION

Gene expression regulation enables differential decoding of the same genetic material. It has generally been studied as a set of sequential processes, with transcription setting the core pattern of expression and translational and post-translational regulation modulating the final output. This concept of an exclusively linear model for the regulation of genetic information decoding is partly the result of the largely isolated discovery and subsequent study of each regulatory step. This approach been necessary for and successful in providing a deep understanding of the biochemical mechanisms that mediate gene regulation, as well as for defining the types of regulatory mechanisms that exist at each level. Such regulatory mechanisms include, for example, transcriptional interference, in which transcription from one promoter locally represses transcription from another (HAUSLER AND SOMERVILLE 1979; ADHYA AND GOTTESMAN 1982; CULLEN *et al.* 1984; EMERMAN AND TEMIN 1984; PROUDFOOT 1986; HIRSCHMAN *et al.* 1988; CORBIN AND MANIATIS 1989; BOUSSADIA *et al.* 1997; GREGER *et al.* 2000; MARTENS *et al.* 2004; KIM *et al.* 2012; VAN WERVEN *et al.* 2012; KIM *et al.* 2016). Transcriptional repression by this mechanism has been associated with cis-enrichment of inhibitory chromatin marks, typically by production of a noncoding transcript (CARROZZA *et al.* 2005; KIM *et al.* 2012; VAN WERVEN *et al.* 2012; KIM *et al.* 2016). An example of a similarly established translational repression mechanism is based on translation of upstream ORFs (uORFs) in the 5’ leader region of some mRNAs at the expense of ORF translation [reviewed in (WETHMAR *et al.* 2010; BARBOSA *et al.* 2013; WETHMAR 2014; HINNEBUSCH *et al.* 2016)]. Typically, uORF-mediated translational repression is viewed as a switch-like mechanism, where the uORFs prevent translation of the downstream ORF under certain conditions, but this repression can be bypassed under other conditions

Recently, we described a form of gene regulation that relies on the integrated use of transcriptional and translational regulation to repress protein expression (CHEN *et al.* 2017; CHIA *et al.* 2017). In budding yeast meiosis, the amount of protein for the conserved kinetochore protein Ndc80 is determined by an unconventional mechanism in which mRNA production from a more distal *NDC80* promoter inhibits Ndc80 protein production through integration of transcriptional and translational interference: the distal promoter-driven transcript cannot be efficiently translated into protein due to uORF translation and its transcription represses the proximal *NDC80* promoter activity in *cis.* In this manner, production of a 5’-extended mRNA isoform inhibits Ndc80 protein production (CHEN *et al.* 2017; CHIA *et al.* 2017). In the case of *NDC80*, the uORF-mediated repression appears to be constitutive, conditional only on the existence of the 5’-extended transcript, rather than condition-specific like the best studied regulatory uORF cases [(MUELLER AND HINNEBUSCH 1986; VATTEM AND WEK 2004), for example]. As such, the re-expression of Ndc80 protein relies on a developmentally induced switch in promoter usage, from distal to proximal, during meiotic progression. Therefore, the coordinated expression of these two functionally distinct *NDC80* mRNA isoforms achieves temporal control of Ndc80 protein expression during meiotic differentiation.

Improved methodology for genome-wide gene expression measurements have resulted in a more complete characterization of the set of transcripts expressed and regions translated than was previously possible. These studies have provided evidence for translation of thousands of uORFs and widespread existence of alternate transcript isoforms [(INGOLIA *et al.* 2009; INGOLIA *et al.* 2011; BRAR *et al.* 2012; DIEUDONNE *et al.* 2015; FLOOR AND DOUDNA 2016; WANG *et al.* 2016), for example], including during the yeast meiotic program. Despite their prevalence, the significance of both alternative transcript production and uORF translation to gene expression output in most of these newly identified cases has been unclear. Using analyses of parallel global mRNA, translation, and protein datasets, we found that many of the uORFs and alternate transcripts seen during the yeast meiotic program were indicative of the mode of coordinate regulation seen for *NDC80*, with at least 379 other genes showing protein levels that are driven by this type of integrated transcriptional and translational control over time through meiotic development (CHENG *et al.* 2018). It was also recently found that this mode of regulation functions to mediate down-regulation of proteins involved in aerobic respiration as a core part of the unfolded protein response [UPR; (Van Dalfsen et al., 2018)]. While these studies were exclusively performed in yeast, we noted that some of the hallmarks of the integrated mode of gene repression that were seen for *NDC80* regulation, and thus used to annotate new cases in yeast, are also known to be common in mammals. For example, almost half of human genes show evidence of alternative promoter usage, resulting in transcript isoforms that differ in their 5’ leader (WANG *et al.* 2016). Additionally, transcripts with extended 5’ leaders that contain uORFs result, in some cases, in a poorly translated transcript compared to isoforms with shorter 5’ leaders (LAW *et al.* 2005; FLOOR AND DOUDNA 2016). Alternative uORF-containing transcripts were also previously defined for several individual mammalian genes, including Mouse double-minute 2 homolog (*MDM2*), an oncogene and repressor of the tumor suppressor p53 (ARRICK *et al.* 1994; BARAK *et al.* 1994; BROWN *et al.* 1999; HUGHES AND BRADY 2005). The *MDM2* transcript isoform produced from the distal P1 promoter contains a longer 5’ leader than the one produced from the proximal P2 promoter (Figure 1B) and this P1-derived *MDM2* isoform specifically is poorly translated due to the presence of two uORFs in its extended 5’ leader, as established by polysome analyses and reporter assays (LANDERS *et al.* 1997; BROWN *et al.* 1999; JIN *et al.* 2003).

**Figure 1.**
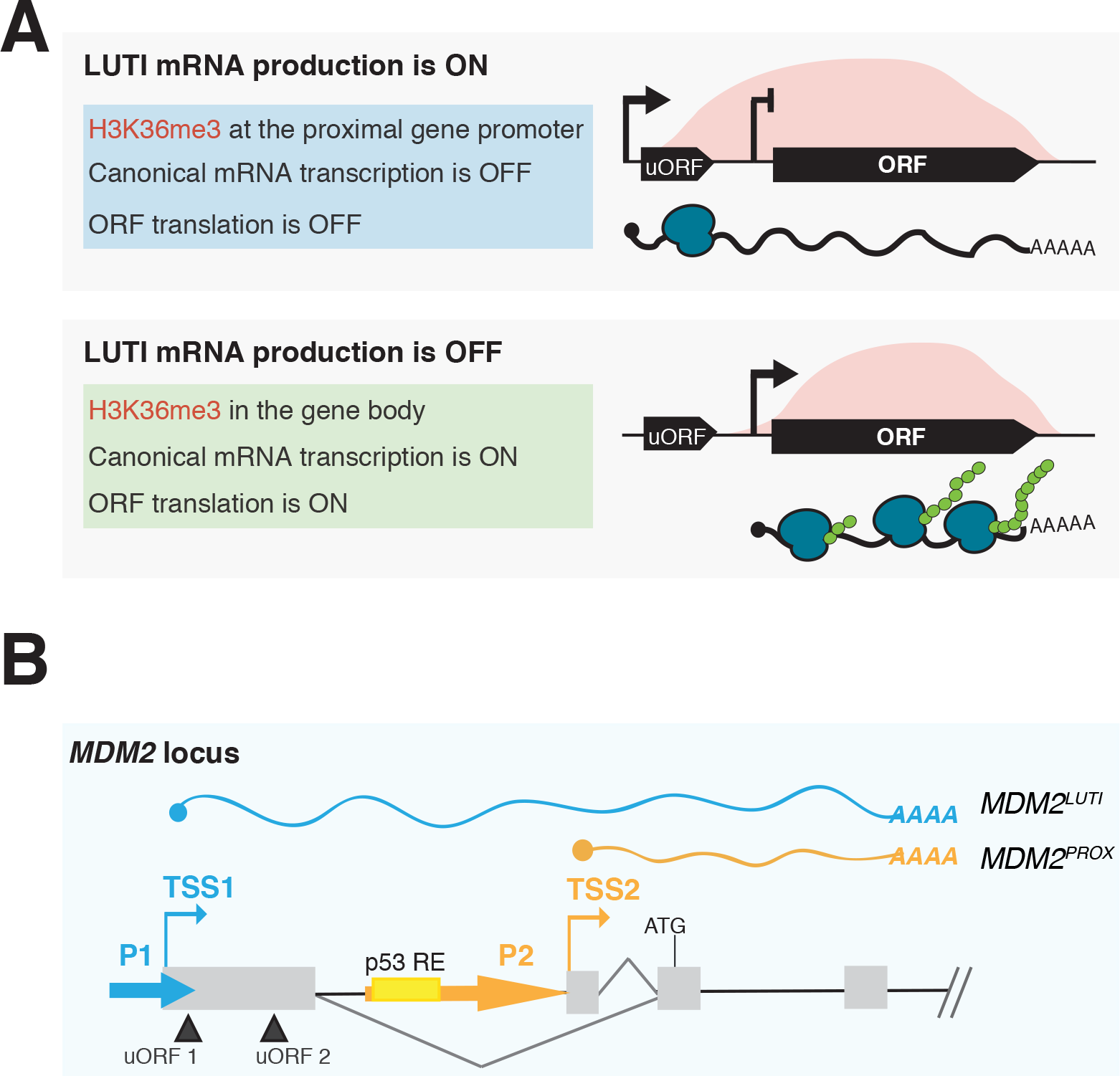
Illustrations of LUTI-based gene repression and *MDM2* gene locus. **A.** Model for LUTI-based gene repression. Top panel: LUTI mRNA production causes an increase in the co-transcriptional H3K36me3 marks at the proximal gene promoter and transcriptional repression of the canonical mRNA isoform. Because LUTI mRNA is not well translated due to uORFs in its extended 5’ leader and because the well-translated canonical mRNA is repressed, the net effect of LUTI mRNA production is the downregulation of translation from the LUTI target gene locus. Bottom panel: In the absence of LUTI expression, transcription from the canonical gene promoter occurs, leading to translation. **B.** Illustration of the *MDM2* gene structure. *MDM2* is transcribed from two different transcription start sites (TSS1 and TSS2) regulated by two different promoters (P1 and P2). Transcription from the distal TSS1 produces a 5’-extended, uORF-containing transcript, which is poorly translated. Hereafter, the P1 promoter-driven transcript isoform is referred to as *MDM2*^*LUTI*^, while the P2-driven isoform, transcribed from the proximal TSS2 is referred to as *MDM2*^*PROX*^. The arrows describe the location of the isoform-specific primers used for the RT-qPCR analyses in this figure and all the subsequent figures (blue arrows: *MDM2*^*LUTI*^ specific primers; yellow arrows: *MDM2*^*PROX*^ specific primers). p53 RE refers to the location of p53 response element within the P2 promoter.

**Table 1.**
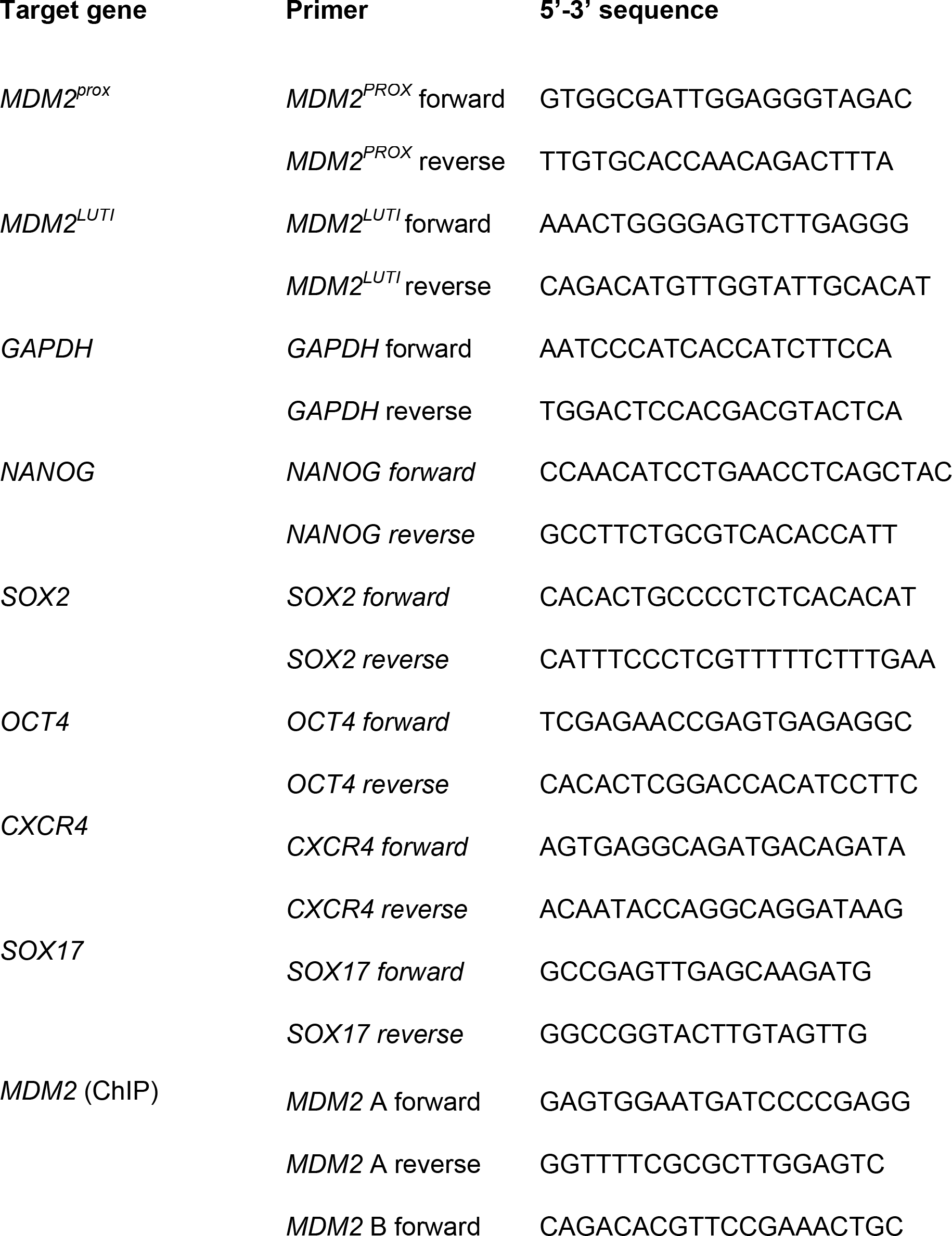
Primers used in this study.

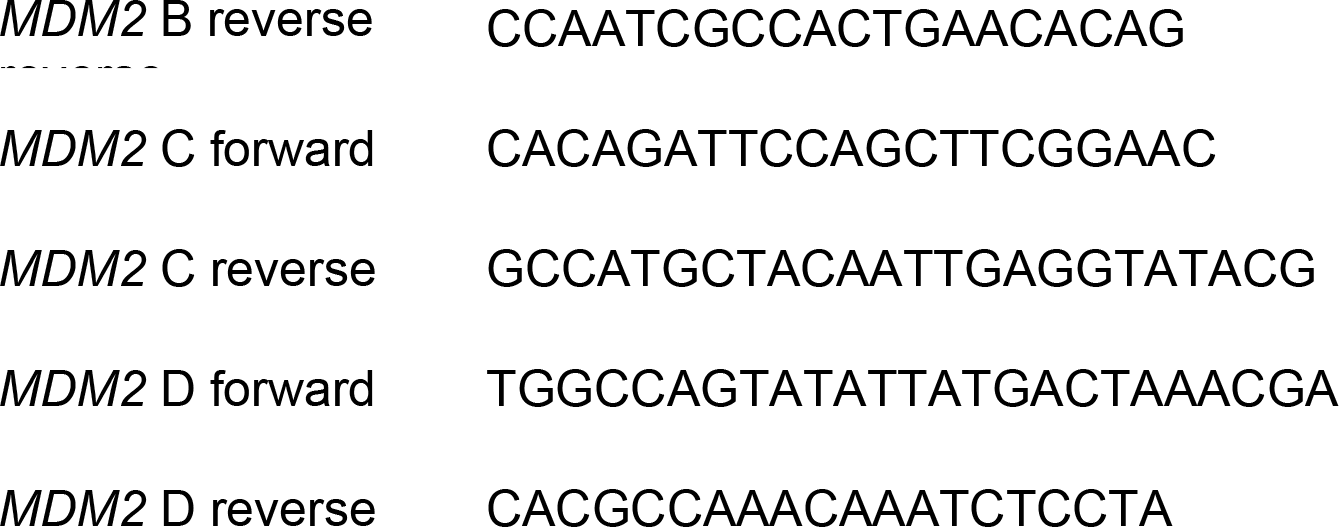

The integrated mode of repression seen for *NDC80* in yeast relies on three key features [Figure 1A, (CHEN *et al.* 2017; CHIA *et al.* 2017)]. First, a developmentally regulated switch between alternative promoters for the same gene leads to the usage of different transcription start sites (TSSs). Second, due to upstream open reading frame (uORF)-mediated translational repression, the distal promoter-generated transcript is inefficiently translated. Third, transcription from the distal promoter represses the expression of the canonical mRNA isoform through transcriptional interference associated with co-transcriptional histone modifications. When all three features exist together, and only then, the activation of *NDC80* transcription can result in a *decrease* in translation from this locus. We termed the distal promoter-generated transcript “*NDC80*^*LUTI*^” for <u>l</u>ong <u>u</u>ndecoded <u>t</u>ranscript <u>i</u>soform, because, despite containing the entire *NDC80* ORF, *NDC80*^*LUTI*^ is not efficiently translated into protein (CHEN *et al.* 2017; CHIA *et al.* 2017). We noted that the regulation at the *MDM2* locus shared features with this LUTI-based mechanism, with the important exception that it was not clear if there was transcriptional interference between the two *MDM2* promoters.

It is well established that *MDM2* P2 can be activated by p53 (WU *et al.* 1993; BARAK *et al.* 1994; HONDA *et al.* 1997), but little is known about the factors that activate P1 and whether transcription from the P1 promoter affects P2 activity had not been explored. To our knowledge, this was true of all other individually and globally defined examples of alternative transcripts of differential translatability, as transcriptional interference and uORF-based translational control have been topics studied independently, and typically by different labs. Transcriptional interference is an essential feature of the LUTI mechanism, because it causes a toggle between the two transcript isoforms, which is necessary for effective gene expression repression. Transcriptional interference also enables efficient developmental regulation in the case of *NDC80*. Evidence of transcriptional interference from nearby transcription has been previously been established in human cells (reviewed in (SHEARWIN *et al.* 2005; PALMER *et al.* 2011), but these cases have not involved production of ORF-encoding mRNAs. Nevertheless, we hypothesized that the type of integrated regulation that we described to be common in yeast might occur at the *MDM2* locus. Here, we report evidence that this is indeed the case, based on observed transcriptional interference between the two promoters at this locus. We also observe developmental regulation of the two *MDM2* transcript isoforms and conclude that the type of integrated transcriptional and translational regulation that we described as a developmental gene regulatory strategy in yeast is also seen in human cells. These findings suggest value in considering translational and transcriptional regulation not only as independent steps, but rather, as potential collaborators in gene expression outcomes.

## RESULTS

A key prediction, if *MDM2* were regulated by a LUTI-based mechanism, would be an inverse relationship between the two *MDM2* transcript isoforms, such that reduction in transcription from P1 should lead to increased transcription from P2. We were able to reliably assay the levels of the P1-and P2-derived transcripts by Real-Time quantitative PCR (RT-qPCR) owing to differential splicing of the two transcripts that results in unique 5’ sequences for both P1 and P2 (Figure S1). To directly test this prediction, we inhibited transcription from P1 by using CRISPRi (GILBERT *et al.* 2013; QI *et al.* 2013). We first examined MCF-7 breast cancer cells stably encoding the catalytically dead Cas9 (dCas9), which is thought to interfere with PolII elongation when targeted near transcription start sites (LARSON *et al.* 2013; QI *et al.* 2013). Expressing each of four different single guide RNAs (sgRNAs) targeting the P1 promoter region led to modest but significant increases, of up to 2-fold, in the P2-derived *MDM2*^*PROX*^ transcript levels, which was associated with the expected reduction of transcription from P1 (Figure 2A and Figure S2A). This result was notable, given that the maximal knockdown of P1 activity was only 40% relative to control cells in these lines (Figure 2A). We tried to enhance the P1 transcriptional knockdown by using CRISPRi in MCF-7 cells that carry a version of dCas9 fused to the Krüppel-associated box (KRAB) transcriptional repression domain (GILBERT *et al.* 2013). However, targeting of dCas9-KRAB to the P1 promoter led to repression of both the P1 and P2 promoters (Figure S3). This finding is consistent with the long-range effect of the KRAB domain up to 1 Kb (GILBERT *et al.* 2014), beyond the 845 bp distance between the P1 and P2 regulated transcription start sites. Therefore, we performed all subsequent experiments using cell lines that stably expressed dCas9 without the KRAB domain, as this first-generation version of CRISPRi allowed us to achieve promoter-specific repression.

**Figure 2.**
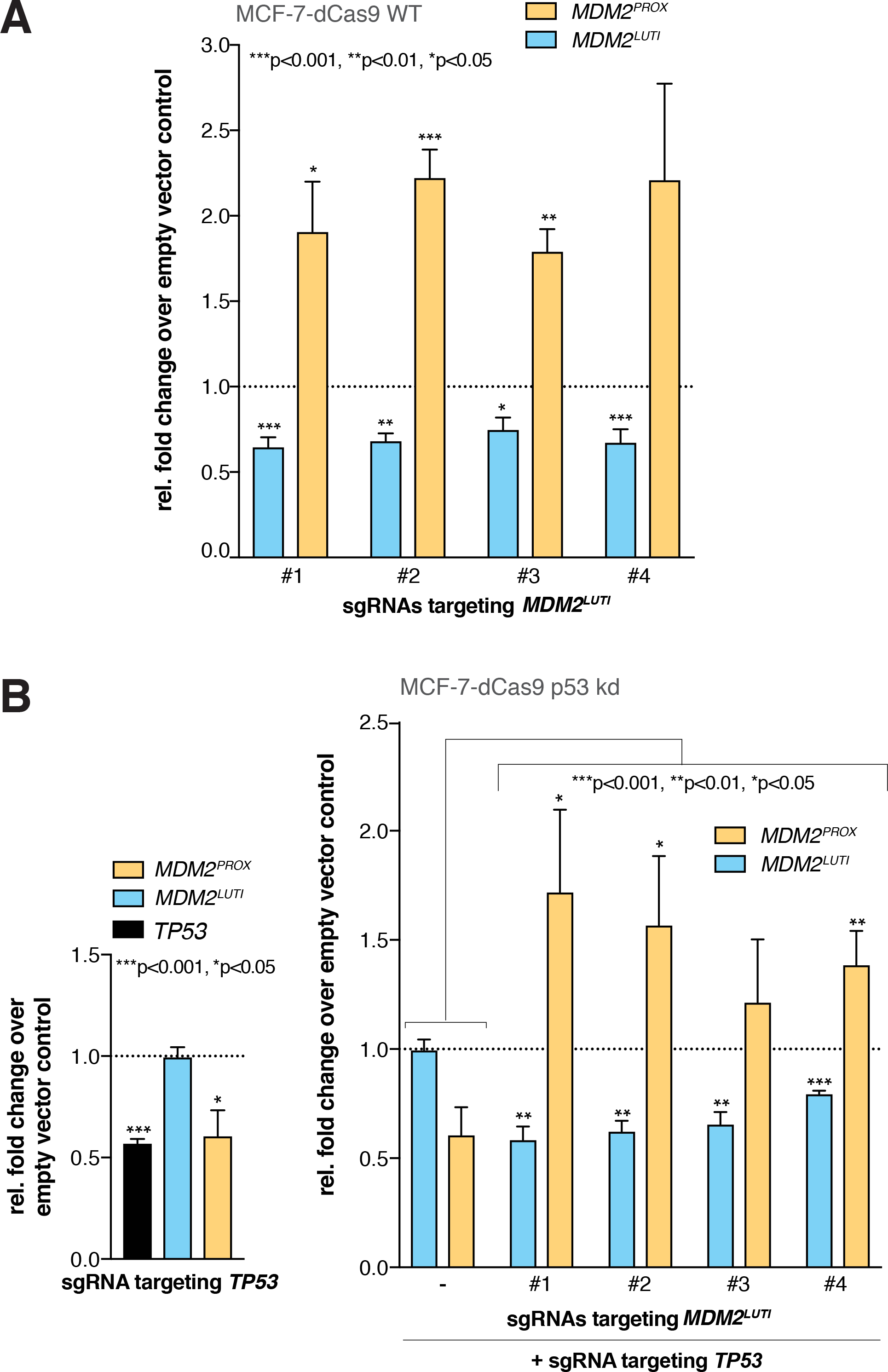
Downregulation of *MDM2*^*LUTI*^ leads to an increase in the expression of *MDM2*^*PROX*^ in MCF-7 cells, independent of p53 expression. **A.** RT-qPCR data displaying the changes of *MDM2*^*LUTI*^ and *MDM2*^*PROX*^ mRNA expression in MCF-7-dCas9 stable cells. The transcription of *MDM2*^*LUTI*^ was inhibited by CRISPRi using four different sgRNAs (#1-4). Data were normalized to *GAPDH*, and the fold change relative to the expression of *MDM2*^*LUTI*^ and *MDM2*^*PROX*^ in the cells transfected with an empty vector was calculated. Data points represent the mean of at least 3 independent biological replicates. Error bars represent standard error of the mean (SEM). Two-tailed Student’s t-test was used to calculate the P-values in this figure and all of the subsequent figures. **B.** RT-qPCR data showing the change in the expression level of *TP53*, *MDM2*^*LUTI*^ and *MDM2*^*PROX*^ in MCF-7-dCas9 cells after CRISPRi-mediated *TP53* knockdown (left) or CRISPRi-mediated p53– and *MDM2^LUTI^-* double knockdown (right, sgRNA #1 through #4), relative to the cells transfected with an empty vector. Data were normalized to *GAPDH.* Data points represent the mean of four biological replicates. Error bars represent SEM.

We further probed the relationship between P1 and P2 by knockdown of the gene encoding p53 (*TP53*) in MCF-7 cells using CRISPRi. Given that p53 is a well-characterized transcriptional activator for P2, it was not surprising that *TP53* knockdown resulted in a significant, 43% reduction of the P2-derived *MDM2*^*PROX*^ transcript (Figure 2B, left panel). However, additional CRISPRi knockdown of the P1-derived transcript, hereon referred to as *MDM2*^*LUTI*^, still resulted in the transcriptional activation of P2, as evidenced by the 2-to 3-fold increase in *MDM2*^*PROX*^ levels in this background compared to the *TP53* knockdown alone (Figure 2B, right panel; Figure S2B; Figure S4). The observation that *MDM2*^*LUTI*^ repression leads to an increase in *MDM2*^*PROX*^ expression, even in cells with reduced p53 levels, suggests that transcription from P1 actively represses P2 activity and that relief of this repression alone can lead to increased expression of *MDM2*^*PROX*^ independent of p53, consistent with transcriptional interference at this locus.

To test whether transcriptional interference based on *MDM2*^*LUTI*^ occurs in a different cell type, we performed similar experiments in K562, a *TP531*^−/−^ myeloid leukemia cell line that routinely shows robust CRISPRi-based repression (GILBERT *et al.* 2014). Inhibition of *MDM2*^*LUTI*^ transcription in these cells resulted in a dramatic increase (up to 10-fold) in *MDM2*^*PROX*^ expression (Figure 3A and Figure S2C). A range of *MDM2*^*LUTI*^ knockdown efficiencies were achieved in this cell line. Consequently, the degree of P1 down-regulation generally correlated with the degree of P2 activation (Figure 3A), suggesting tunability of the transcriptional interference at this locus.

**Figure 3.**
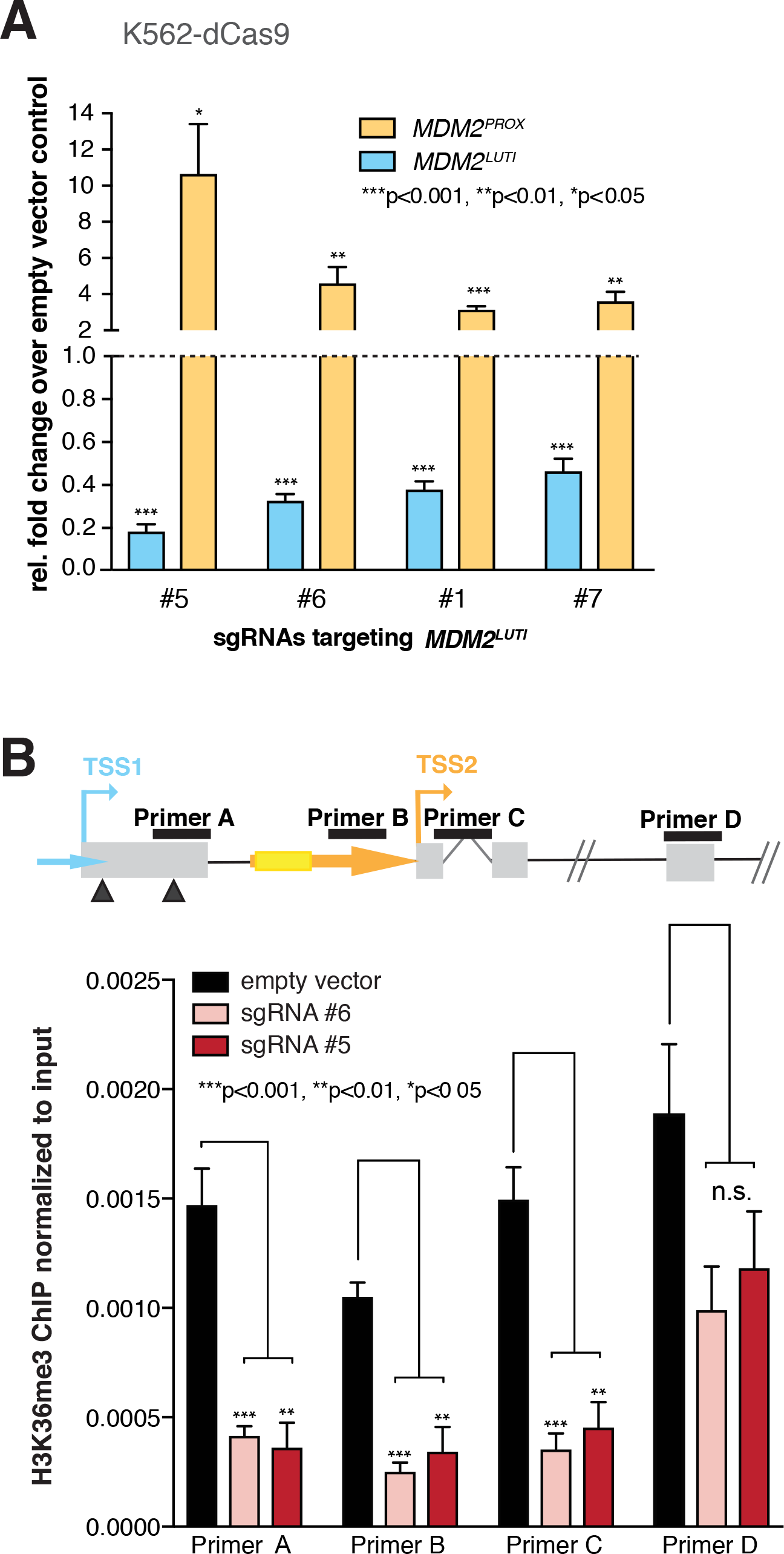
Downregulation of *MDM2*^*LUTI*^ reduces repression-associated histone marks at the P2 promoter. **A.** RT-qPCR data displaying the changes in *MDM2*^*LUTI*^ and *MDM2*^Prox^expression levels in a stable K562-dCas9 cell line in which the transcription of *MDM2*^*LUTI*^ had been inhibited by CRISPRi using four different sgRNAs (#5, #6, #1, and #7). Data were normalized to *GAPDH*, and the fold change relative to cells transfected with the empty vector was calculated. Data points represent the mean of at least 3 independent biological replicates. Error bars represent SEM. **B.** Chromatin immunoprecipitation (ChIP) data displaying H3K36 trimethylation (H3K36me3) enrichment around the proximal TSS (TSS2) in K562-dCas9 cells after CRISPRi-mediated inhibition of *MDM2*^*LUTI*^ expression. Location of the four different primer pairs (A, B, C, and D) are shown in the schematic above the graph Arrowheads indicate the location of uORFs in exon 1 and yellow box indicates the p53 response element. Data points represent the mean of 4 independent biological replicates. Error bars represent SEM. n.s. = not significant.

H3K36me3 is a co-transcriptionally established modification that occurs in regions downstream of active promoters (XIAO *et al.* 2003; BANNISTER *et al.* 2005; MIKKELSEN *et al.* 2007), and in budding yeast is associated with a decrease in spurious transcription initiation from within transcribed genes (LI *et al.* 2003; XIAO *et al.* 2003; CARROZZA *et al.* 2005; KEOGH *et al.* 2005; KIM *et al.* 2016) and in noncoding RNA transcription-dependent repression of gene promoters (KIM *et al.* 2012; VAN WERVEN *et al.* 2012). H3K36me3 is enriched at the proximal *NDC80* promoter as a result of *NDC80*^*LUTI*^ transcription and is involved in the transcriptional interference seen at the *NDC80* locus (CHIA *et al.* 2017). In mammalian cells, H3K36me3 has been implicated in silencing, including the repression of spurious intragenic transcription [(DHAYALAN *et al.* 2010; XIE *et al.* 2011; WAGNER AND CARPENTER 2012; BAUBEC *et al.* 2015; SUZUKI *et al.* 2017), but its involvement in promoter repression has been less clear. We found that down-regulation of *MDM2*^*LUTI*^ expression resulted in a greater than 3-fold decrease in the H3K36me3 signal over the P2 promoter (Figure 3B, Figure S5). In contrast, H3K36me3 signal remained high within the *MDM2* gene body, likely due to increased *MDM2*^*PROX*^ transcription under these conditions. These data are consistent with a mechanism whereby *MDM2*^*LUTI*^ expression represses transcription from the P2 promoter through co-transcriptional histone modifications, and provide further support for a model in which the proximal *MDM2* promoter is controlled by a similar mechanism to that defined in yeast.

In budding yeast, developmentally controlled switching between the LUTI and canonical mRNA isoforms occurs pervasively during meiotic differentiation (CHENG *et al.* 2018). To begin to test whether such transcript toggling naturally occurs in human cells, we used two different human Embryonic Stem Cell (hESC) differentiation models. In human hESCs, both *MDM2* transcript isoforms were expressed (Figure 4A). When these cells were induced to undergo neuronal differentiation, a transient switch in transcript isoform expression from *MDM2*^*PROX*^ to *MDM2*^*LUTI*^ was evident between hESCs and neuronal precursors (NPCs) (Figure 4B). We also observed an anti-correlation between *MDM2*^*PROX*^ and *MDM2*^*LUTI*^ expression as hESCs differentiated into an endodermal fate, as determined by endoderm-specific markers (Figure 4C, Figure S6). This inverse pattern of proximal and distal promoter usage seen during hESC differentiation suggests that the LUTI-based mechanism regulates *MDM2* expression during normal cellular differentiation.

**Figure 4.**
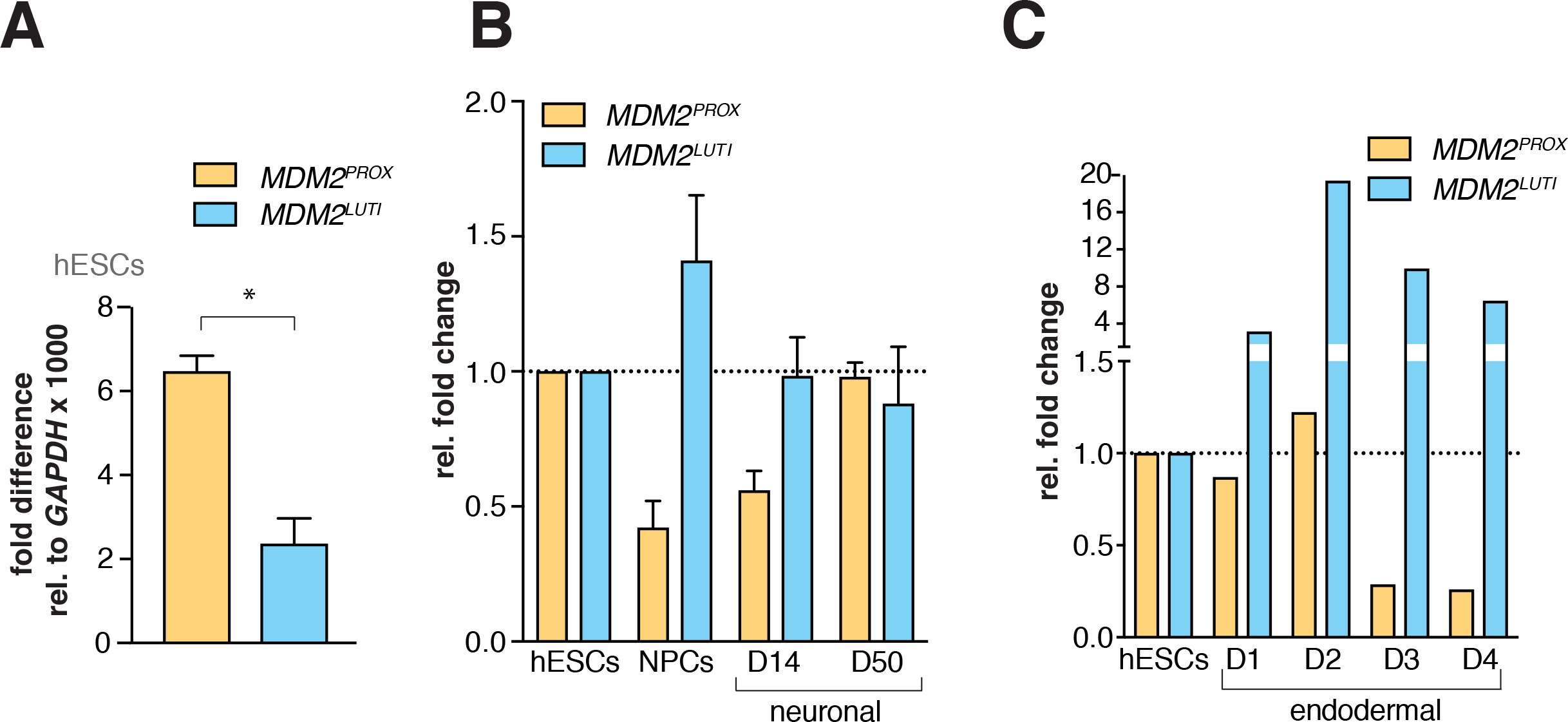
P1 and P2-driven *MDM2* transcript isoform toggling can be seen during human embryonic stem cell differentiation. **A.** RT-qPCR data showing the fold differences of *MDM2*^*LUTI*^ and *MDM2*^*PROX*^ transcript levels relative to *GAPDH* expression levels x1000 in human embryonic stem cells (hESCs). Error bars represent the range measured for two biological replicates. **B.** RT-qPCR data sowing the relative expression of *MDM2*^*LUTI*^ and *MDM2*^*PROX*^ transcripts in human embryonic stem cells (hESCs), neural progenitor cells (NPCs), Day 14 and Day 50 neurons (D14 and D50). Data were normalized relative to *MDM2*^*LUTI*^ or *MDM2*^*PROX*^ transcript abundance in hESCs. Error bars refer to the range measured for two biological replicates. **C.** RT-qPCR data showing the changes in the expression of *MDM2*^*PROX*^ and *MDM2*^*LUTI*^ in hESCs differentiating into endoderm. D1-D4 refers to days after transfer of the hESCs to endoderm differentiation medium. Data were normalized relative to *MDM2*^*LUTI*^ or *MDM2*^*PROX*^ transcript abundance in hESCs.

## DISCUSSION

We report here that an integrated transcriptional and translational regulatory strategy, initially defined for *NDC80* in yeast, also occurs in humans. Based on the ubiquitous use of alternative promoters and uORF translation in humans (INGOLIA *et al.* 2011; FLOORAND DOUDNA 2016; WANG *et al.* 2016; TRESENRIDER AND UNAL 2018), this type of regulation may control the expression of many mammalian genes. Both the type of transcriptional control described here and the uORF-based translational regulation already established for the distal promoter-derived *MDM2* transcript are known modes of gene regulatory control. Our study shows that transcriptional interference can be *integrated* with production of alternate transcripts of differential translatability in mammals, in a manner that resembles the LUTI-based regulation seen in yeast. These findings suggest value in revisiting the linear model of gene expression control that dominates the interpretation of past and current data in favor of a more holistic view of the interplay between different levels of gene expression control.

Canonical models to explain the prevalence of mammalian alternative promoter usage, for example, suggest that one promoter might serve as a “back-up” or that the use of two promoters could simply allow activation by different transcription factors that are present in different cell types (DAVULURI *et al.* 2008). However, in the case of *MDM2*, we argue that its two promoters are fundamentally different in function. The P1 promoter produces a poorly translated *MDM2*^*LUTI*^ transcript and the production of *MDM2*^*LUTI*^ from this promoter interferes with P2 activity in *cis*, reducing the transcription of the well-translated *MDM2*^*PROX*^ isoform. Therefore, P1-driven *MDM2*^*LUTI*^ mRNA production serves to downregulate *MDM2* expression. Further, this repression occurs in a tunable manner. (Figure 5). A similar trend was also observed for the *Adh* gene in fruit fly, which is expressed from two distinct promoters (JORGENSEN *et al.* in preparation). Both findings are consistent with the notion that transcriptional interference can be used to tune gene expression, rather than acting as an on-off switch. (CHIA *et al.* 2017).

**Figure 5.**
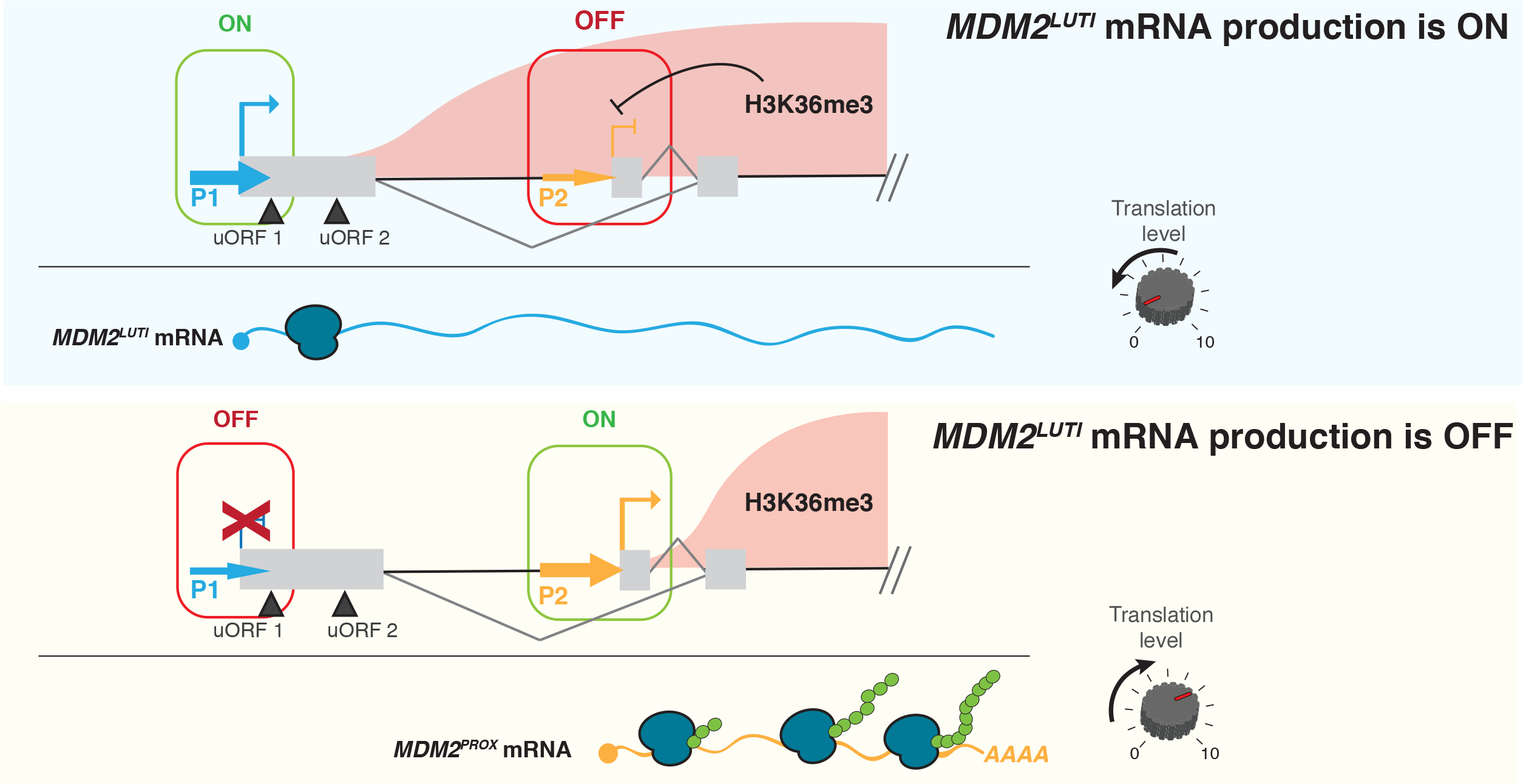
Model of the LUTI mRNA based mechanism of the *MDM2* gene. The *MDM2* gene has two promoters, P1 and P2. *MDM2*^*PROX*^ is regulated by P2 whereas *MDM2*^*LUTI*^ is regulated by P1. In comparison to *MDM2*^*PROX*^, *MDM2*^*LUTI*^ is poorly translated because of the existence of two upstream open reading frames (uORFs) in its extended 5’-leader. Top panel: When P1 promoter is active (“ON”), *MDM2*^*LUTI*^ transcription establishes H3K36 trimethylation at the downstream P2 promoter and causes repression of P2 (“OFF”). As a result, *MDM2*^*LUTI*^ becomes the predominant transcript product from the *MDM2* locus. Bottom panel: When P1 promoter is “OFF”, transcriptional repression of the downstream P2 promoter is relieved, culminating in the expression of *MDM2*^*PROX*^. *MDM2*^*PROX*^ is efficiently translated, resulting in higher MDM2 translation.

Contrary to traditional gene regulatory models, mRNA and protein abundances generally show poor correlations over developmental programs in genome-scale yeast and vertebrate studies (PESHKIN *et al.* 2015; CHENG *et al.* 2018). In yeast, hundreds of such cases can be explained by LUTI-based regulation, whereby developmentally regulated transcript distal and proximal promoter usage drives final protein output levels (CHENG *et al.* 2018). In these conditions, we found that a single developmentally regulated transcription factor could drive distinct sets of canonical and LUTI targets, which resulted in coordinate up-and down-regulation of protein levels for the two target sets (CHENG *et al.* 2018). We propose that this mechanism may be advantageous for developmental programs, which are typically characterized by multiple transcription-factor driven switches in cell stage, because of the ability to temporally coordinate up-and down-regulation of gene sets at each transition. In principle, whether a gene is in one set or another could be defined simply by the position of the binding site for a given transcription factor relative to the ORF and uORF sequences (CHIA *et al.* 2017; CHENG *et al.* 2018; OTTO AND BRAR 2018; TRESENRIDER AND UNAL 2018). The natural toggling between *MDM2* isoforms during differentiation shown here suggests that the broad use of this type of regulation for developmental modulation of gene expression may be conserved.

MDM2 levels are elevated in a variety of cancers (reviewed in (RAYBURN *et al.* 2005) and this elevation has been attributed in some cases to an increase in translation of the pool of *MDM2* transcripts, based on increased transcription from the P2 (LANDERS *et al.* 1997; CAPOULADE *et al.* 1998; BROWN *et al.* 1999). Much research has focused on identifying alternate transcription factors that can activate P2—as it is clear that transcription can occur from this promoter in the absence of p53—and several have been found (PHELPS *et al.* 2003; ZHANG *et al.* 2012), but relatively little is known about P1 regulation. Our study argues that MDM2 expression levels could be modulated by changes in activation of P1 alone, suggesting a promising new general area for the development of gene regulatory tools that modulate P1 activity, and the activity of other yet-to-be-identified LUTI mRNA promoters, as a means to fine tune gene expression.

## MATERIALS AND METHODS

### Cell lines

MCF-7-dCas9 and -dCas9-KRAB cells were cultivated at 37°C with 5% CO_2_ in high glucose Dulbecco’s modified Eagle media (GlutaMax, Gibco) supplemented with 10% FBS and 1% penicillin-streptomycin. K562-dCas9 cell lines were cultivated at 37°C with 5% CO_2_ in RPMI1640 media (Gibco) supplemented with 10% FBS and 10mM HEPES. hESCs (WIBR3 NIH#0079) were maintained in culture as described in (LENGNER *et al.* 2010). The differentiation into definitive endoderm was performed using the STEMdiff™ Definitive Endoderm Kit (Stem Cell Technologies) following the manufacturer’s instructions. MCF-7-dCas9 and -dCas9-KRAB cells were kindly provided by Howard Y. Chang (Stanford University). K562-dCas9 cells were kindly provided by Jonathan Weissman (University of California, San Francisco). Cell lines were authenticated by STR profiling and were tested to be negative for mycoplasma (MycoAlert™ Mycoplasma Detection Kit, Lonza).

### RNA isolation, cDNA synthesis and quantitative polymerase chain reaction

Total RNA from hESCs differentiating into definitive endoderm and hESCs differentiating into neurons, MCF-7-dCas9, MCF-7-dCas9-KRAB and K562-dCas9 cells was isolated using Trizol (Life Technologies) according to the manufacturer’s instructions. Equal amounts of RNA were primed with random hexamers and reverse transcribed using SuperScript II Reverse Transcriptase (Thermo Fisher) according to the manufacturer’s instructions. RNA levels were quantified using SYBR Green/Rox (ThermoFisher) and the StepOnePlus Real-time PCR system (ThermoFisher).

Samples for total RNA isolation from hESCs differentiating into neurons (hESC, NPC, neurons D14, neurons D50) were a gift from Helen Bateup [University of California, Berkeley]

### CRISPRi knockdowns

SgRANs targeting *MDM2*^*LUTI*^ were designed and cloned into the lentiviral pU6-sgRNA EF1Alpha-puro-T2A-BFP vector. Lentivirus was packaged by co-transfecting sgRNA-expression plasmids and the packaging vectors pCMV-dR8.91 and pMD2.G into 293T cells using the TransIT^®^-LT1 Transfection Reagent (Mirus). Cells were treated with ViralBoost (Alstem) to allow for efficient lentivirus production and lentivirus was harvested 72 hours post-transfection. CRISPRi-directed gene knockdown was achieved by transducing MCF-7-dCas9, -dCAS9-KRAB and K562-dCas9 cell lines with sgRNA-containing lentivirus in the presence of 8μg/ml polybrene (Millipore Sigma). Successfully transduced cells were puromycin-selected (ThermoFisher; 13μg/ml for MCF-7 and 3μg/ml for K562 cells) and harvested 7 days post-infection. The pU6-sgRNA EF1Alpha-puro-T2A-BFP vector was a gift from Jonathan Weissman (Addgene plasmid # 60955).

### H3K36me3 Chromatin immunoprecipitation (ChIP)

K562-dCas9 cells (5 × 15cm^2^ plates per sample) were treated with 1% formaldehyde 16%, methanol free, Ultra Pure, Polysciences) for 10 minutes at room temperature to crosslink DNA and protein. The crosslinking reaction was stopped by adding 0.125M PBS-glycine and cells were harvested by centrifugation. Cells were subsequently resuspended in ice-cold PBS containing 0.25mM PMSF and 10ug/ml aprotinin (Millipore) and pelleted by centrifugation. Chromatin immunoprecipitation of these pellets was performed as previously described (TESTA *et al.* 2005) with minor modifications. Chromatin was sonicated 50 × 30 seconds ON/30 seconds OFF with a Bioruptor^®^ Pico (Diagenode) to obtain fragment sizes of ~200 bp. The sheared samples were incubated in RIPA buffer II (10 mM Tris-Cl, pH 8.0, 1 mM EDTA, pH 8.0, 0.5 mM EGTA, 1% Triton X-100, 0.1% SDS, 0.1% Na-deoxycholate, 140 mM NaCl) containing protease inhibitors and PMSF, with Dynabeads^®^ Protein A (Invitrogen) for 2 h at 4 °C on rotation. After removal of Dynabeads^®^ Protein A, precleared lysates were immunoprecipitated overnight with 4 ug of rabbit anti-mouse IgG (Ab46540, Abcam) or anti-Histone H3 tri methyl lysine 36 (Ab9050, Abcam). Immunoprecipitates were recovered by incubation for 2 h at 4 °C with previously blocked Protein A Dynabeads in RIPA buffer II (1 μg/μl bovine serum albumin, protease inhibitors, and PMSF). Reverse crosslinked input DNA and immunoprecipitated DNA fragments were amplified with SYBR Green/Rox (ThermoFisher) and quantified with StepOnePlus Real-time PCR system (ThermoFisher).

## Data availability

All the reagents generated in this study are available upon request.

## Acknowledgements

We are grateful to Howard Chang and Sueng Woo Cho for providing dCas9 and dCas9-KRAB MCF-7 cell lines; Luke Gilbert, Max Horlbeck, and Jonathan Weissman for providing K562 dCas9 cell lines and for extensive technical help; Helen Bateup, John Blair and Dirk Hockemeyer for providing hESC differentiation samples; Stephen Floor for providing advice and sharing data prior to publication and Christiane Brune for technical support. We thank Barbara Meyer, Jasper Rine, Michael Rape, Folkert van Werven, Jingxun Chen and Eric Sawyer for their valuable feedback on the manuscript; and members of the Tjian and Darzacq labs for tissue culture assistance and advice. This work was supported by funds from the Pew Charitable Trusts (00027344), Damon Runyon Cancer Research Foundation (35-15), National Institute of Health (DP2 AG055946-01) and Glenn Foundation for Medical Research to E.Ü.; funds from the Pew Charitable Trusts (00029624), the Alfred P. Sloan Foundation (FG-2016-6229), and the National Institute of Health (DP2 GM-119138) to G.B.; funds from the California Institute of Regenerative Medicine (CIRM, LA1-08013) to X.D., funds from the Howard Hughes Medical Institute (003061) to R.T. and funds from the Shurl and Kay Curci Foundation to L.Y.C.

## Author contributions

IH, AT, XD, RT, LYC, EÜ, and GAB designed experimental strategies and methods and performed data analysis; IH, JCB, VJ, AT, CDD, and LYC executed experiments; IH, EÜ, and GAB wrote the manuscript.

## SUPPLEMENTAL FIGURE LEGENDS

**Figure S1. Information about the location of the single guide RNAs (sgRNAs) used for the *MDM2* locus in this study**. Binding sites of the sgRNAs (red lines) used for the CRISPRi-mediated knockdown of *MDM2*^*LUTI*^ within the *MDM2* gene. Locations of primers used to amplify *MDM2*^*LUTI*^ (green) and *MDM2*^*PROX*^*(purple)* in RT-qPCRs.

**Figure S2. Downregulation of *MDM2*^*LUTI*^ expression increases levels of *MDM2*^*PROX*^**.

RT-qPCR data showing changes in *MDM2*^*PROX*^ and *MDM2*^*LUTI*^ expression upon CRISPRi-mediated knockdown of *MDM2*^*LUTI*^ relative to *GAPDH* expression levels x1000 in MCF-7-dCas9 WT cells (**A**), MCF-7-dCas9 cells in which the expression of *TP53* was downregulated (**B**), and K562-dCas9 cells (**C**). Error bars represent the SEM of at least three independent biological replicates.

**Figure S3. Both *MDM2*^*PROX*^ and *MDM2*^*LUTI*^ levels are reduced upon CRISPRi targeting of *MDM2*^*LUTI*^ transcription start site in MCF-7-dCas9-KRAB cell lines**. RT-qPCR data showing the changes in *MDM2*^*PROX*^ and *MDM2*^*LUTI*^ expression upon CRISPRi-mediated knockdown of *MDM2*^*LUTI*^ in MCF-7-dCas9 (solid bars) and MCF-7-dCas9-KRAB cell lines (checkered bars), using two different sgRNAs (#1 and #3). Data were normalized to *GAPDH* and the fold change relative to cells transfected with the empty vector was calculated.

**Figure S4. *MDM2*^*PROX*^ can be up-regulated upon *MDM2*^*LUTI*^ downregulation, even under conditions with low p53 levels**. RT-qPCR data showing the expression of *MDM2*^*PROX*^ and *MDM2*^*LUTI*^ in the MCF-7-dCas9 cells treated with different sgRNAs (#1-4) targeting *MDM2*^*LUTI*^ in the presence (+) or absence (−) of a sgRNA targeting *TP53.* Data were normalized to *GAPDH* and the fold change relative to cells transfected with the empty vector (with our without sgRNA targeting *TP53)* was calculated. Data points represent the mean of at least 3 independent biological replicates. Error bars represent SEM.

**Figure S5. Quality assessment for the H3K36me3 ChIP**. qPCR analysis of chromatin immunoprecipitation performed with IgG or anti-H3K36me3. Same primers sets were used as in Figure 3C. Note that the H3K36me3 data are the same as shown in Figure 3C. Data points represent the mean of 4 independent biological replicates. Error bars represent SEM.

**Figure S6. Validation of endodermal differentiation of human embryonic stem cells**. RT-qPCR data showing the changes in expression of hESC-specific genes (*NANOG*, *SOX2*, and *OCT4*) and endoderm-specific genes (*SOX17* and *CXCR4*) during endodermal differentiation (Endo D1 through D4). Values normalized to the expression in hESCs.

